# A single-domain response regulator functions as integrating hub to coordinate general stress response and development in alpha-proteobacteria

**DOI:** 10.1101/299479

**Authors:** C. Lori, A. Kaczmarczyk, I. de Jong, U. Jenal

## Abstract

The alpha-proteobacterial general stress response is governed by a conserved partner-switching mechanism that is triggered by phosphorylation of the response regulator PhyR. In the model organism *Caulobacter crescentus* PhyR was proposed to be phosphorylated by the histidine kinase PhyK, but biochemical evidence in support of such a role of PhyK is missing. Here, we identify a single-domain response regulator, MrrA, that is essential for general stress response activation in *C. crescentus*. We demonstrate that PhyK does not function as a kinase but accepts phosphoryl groups from MrrA and passes them on to PhyR, thereby adopting the role of a histidine phosphotransferase. MrrA is phosphorylated by at least six histidine kinases that likely serve as stress sensors. MrrA also transfers phosphate to LovK, a histidine kinase involved in *C. crescentus* holdfast production and attachment, that also negatively regulates the general stress response. We show that LovK together with the response regulator LovR acts as a phosphate sink to redirect phosphate flux away from the PhyKR branch. In agreement with the biochemical data, a *mrrA* mutant is unable to activate the general stress response and shows a hyper-attachment phenotype, which is linked to decreased expression of the major holdfast inhibitory protein HfiA. We propose that MrrA serves as a central phosphorylation hub that coordinates the general stress response with *C. crescentus* development and other adaptive behaviors. The characteristic bow-tie architecture of this phosphorylation network with MrrA as the central knot may expedite the evolvability and species-specific niche adaptation of this group of bacteria.

**Importance:** Two-component systems (TCSs) consisting of a histidine kinase and a cognate response regulator are predominant signal transduction systems in bacteria. To avoid cross-talk, TCS are generally thought to be highly insulated from each other. However, this notion is based largely on studies of the HisKA subfamily of histidine kinases, while little information is available for the HWE and HisKA2 subfamilies. The latter have been implicated in the alpha-proteobacterial general stress response. Here, we show that in the model organism *Caulobacter crescentus* an atypical FATGUY-type single-domain response regulator (SDRR), MrrA, is highly promiscuous both in accepting and transferring phosphoryl groups from and to a number of up- and downstream kinases, challenging the current view of strictly insulated TCSs. Instead, we propose that FATGUY response regulators have evolved in alpha-proteobacteria to serve as central phosphorylation hubs to broadly sample information and distribute phosphoryl groups between the general stress response pathway and other TCSs, thereby coordinating multiple cellular behaviors. This signaling cascade includes presumable histidine kinases harboring intact catalytic and ATP-binding (CA) domains that, however, do not function as kinases, but instead have adopted a role as histidine phosphotransferases. Our work highlights a complex phosphorylation network in alpha-proteobacteria that coordinates the general stress response with changes in other cellular behaviors and development.

## Introduction

All living organisms must constantly monitor their environment to ensure survival and successful reproduction. In bacteria, adaptive responses often involve motility or chemotaxis, the formation of surface-grown multicellular biofilms, or general and specific stress responses. Through altered behavior or physiological states bacterial cells can withstand or escape potentially harmful or suboptimal conditions that endanger their fitness. However, adaptive responses interfere with normal development or proliferation and, conversely, specific developmental stages may shape an organism’s ability to tolerate and respond to environmental changes. For instance, in *C. crescentus* the ability to escape or withstand unfavorable conditions changes during the reproductive cycle. *C. crescentus* has a dimorphic lifestyle that, upon division, produces a motile and a sessile daughter (1). The motile swarmer cell (SW) is equipped with a flagellum and is able to perform chemotaxis but remains in a replication incompetent state. To proliferate, the SW cell needs to differentiate into a sessile stalked cell, a process during which it loses its flagellum, synthesizes an adhesin called the holdfast and initiates replication and cell division. Intriguingly, the ability of *C. crescentus* to survive stressful conditions depends on the cell cycle stage (2). Moreover, cells experiencing stress or suboptimal growth conditions respond by adjusting their development and cell cycle progression (3). For instance, when starved for carbon cells respond by blocking cell cycle progression and chromosome replication (4, 5). In contrast, *C. crescentus* cells experiencing heat or ethanol stress respond by over-replicating their chromosomes (5). While these responses are thought to increase bacterial survival, the underlying molecular mechanisms coordinating the stress response with developmental or reproductive processes remain largely unknown.

Bacterial signal transduction is predominated by two-component phosphorylation cascades (6). Generally, a histidine kinase undergoes autophosphorylation on a conserved histidine residue upon perception of a specific external or internal stimulus. The phosphoryl group is then transferred to a conserved aspartate residue of the receiver (Rec) domain of a cognate response regulator. Rec modification in turn controls the activity of various response regulator output domains (7). A subclass of response regulators, called single-domain response regulators (SDRRs), lacks a dedicated output domain comprising only the phosphoryl-accepting Rec domain (8). These proteins are thought to act by directly interacting with other proteins and allosterically modulating their activity (9, 10); or as shuttles or sinks, transferring phosphoryl groups between phosphorelay components or draining phosphate away from response regulators (11–14). The *C. crescentus* genome encodes a total of 20 SDRRs, a large fraction of which interact with the flagellar motor similar to the canonical CheY protein in *E. coli* (15). Two SDRRs, DivK and CpdR, are members of a complex regulatory network controlling the activity of the cell cycle regulator CtrA, a central response regulator mediating *C. crescentus* proliferation and behavior (16–18). While DivK acts as allosteric regulator of several cell cycle kinases that are positioned upstream of CtrA (10, 19–21), CpdR serves as a protease adaptor to control cell cycle-dependent degradation of CtrA (18, 22).

Functional information is available for one additional member of the SDRR family in *C. crescentus*, LovR. LovR is encoded in the *lovKR* operon, with LovK considered to be the cognate kinase of LovR. The LovK/LovR two-component system was proposed to control *C. crescentus* surface attachment in response to blue light by promoting the production of adhesive holdfast (23)(24). More recently, the LovKR proteins were shown to also negatively regulate the general stress response, an adaptive response to a diverse range of adverse environments and important for survival under harmful conditions (25). The alpha-proteobacterial general stress response is conserved in essentially all free-living members of this class and is controlled by a partner-switching mechanism involving the sigma factor SigT (or EcfG), the anti-sigma factor NepR, and the response regulator and anti-anti sigma factor PhyR (26–28) (Figure 1A). In the absence of stress, PhyR is dephosphorylated and NepR interacts with SigT, preventing the sigma factor from productive interaction with RNA polymerase. When cells experience stress, PhyR is phosphorylated and binds NepR, resulting in SigT release and the activation of its target genes. While in *C. crescentus* PhyK was proposed to be the major histidine kinase of PhyR, LovK and LovR were proposed to play a role in PhyR dephosphorylation (25, 29). The mechanistic details of this process have remained elusive.

**Figure 1:**
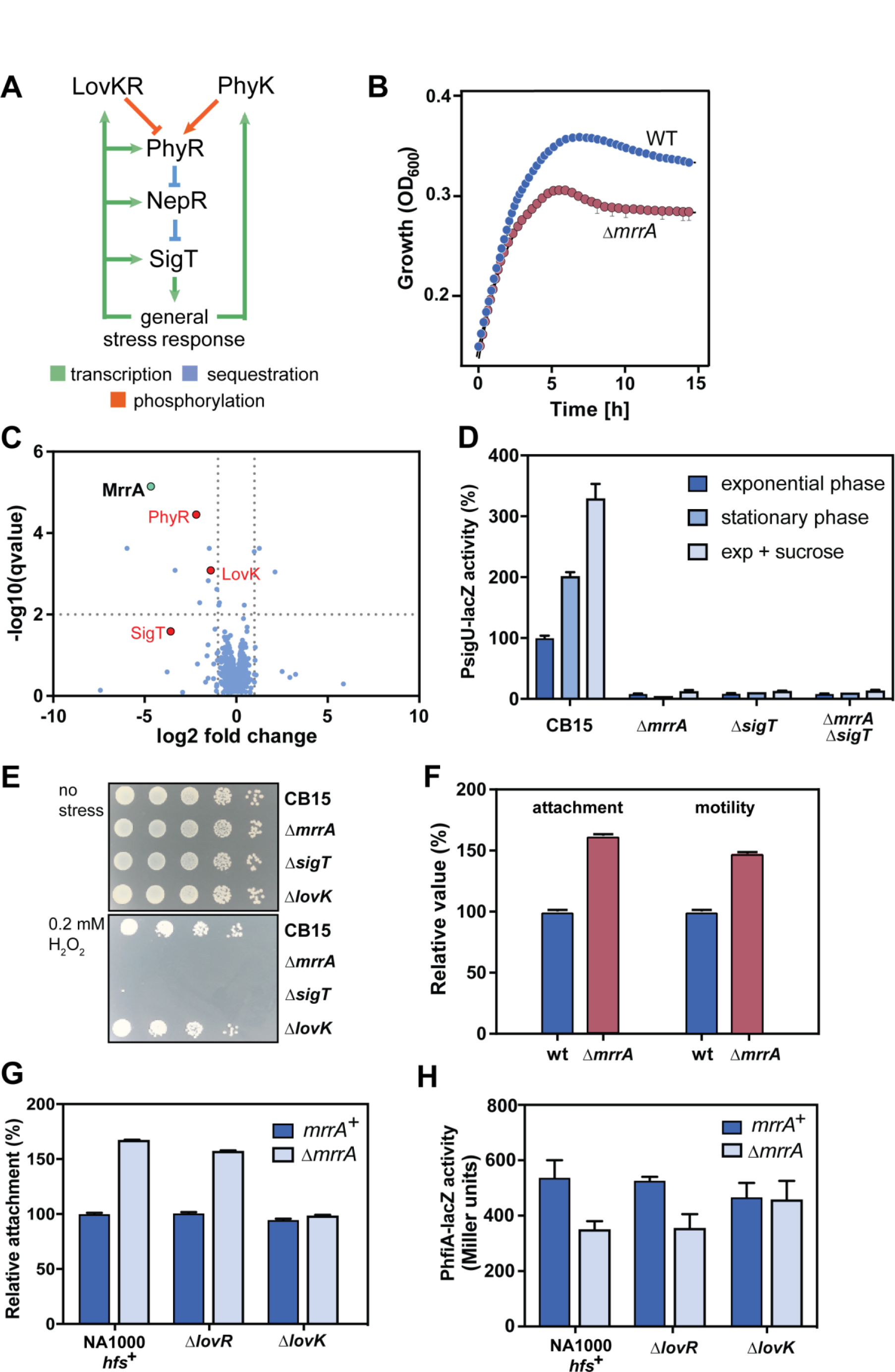
MrrA controls *C. crescentus* behavior and stress response. **(A)** Schematic representation of the general stress response pathway in alpha-proteobacteria. **(B)** A Δ*mrrA* mutant shows decreased growth upon entry into stationary phase. Growth was measured in 96-well plates by periodically monitoring optical density at 660 nm. **(C)** Comparison of proteomes of *C. crescentus* wild type and *ΔmrrA* mutant. On the x-axis log2 fold changes of the *ΔmrrA*/WT ratios are plotted and on the y-axis log10(qvalue). Red dots highlight targets that were significantly downregulated and were suggested to be involved in general stress response. MrrA is highlighted in green. **(D)** MrrA is required for SigT-dependent gene expression. LacZ reporter fusions were used to determine the activity of the SigT-dependent *sigU* promoter. Cells were grown in PYE or PYE supplemented with 150 mM sucrose with two biological and several technical replicates being used per strain and condition. Error bars indicate the standard deviation. **(E)** MrrA is required for efficient survival under stress. Cells were grown to exponential phase in minimal medium with xylose (M2X) and stressed using 0.2 mM H_2_O_2_ for 1 hour. Serial 1:10 dilutions are shown. **(F)** MrrA controls *C. crescentus* surface attachment and motility. Relative values of overall attachment and motility are shown and were determined as outlined in materials and methods. **(G)** LovK is essential for the surface attachment repression of MrrA. Relative values of overall attachment are shown. **(G)** MrrA increases *hfiA* expression in a LovK-dependent manner. Cells harboring *hfiA*-*lacZ* reporter fusions were grown in PYE and β-galactosidase activities determined as described in materials and methods. For all experiments shown in this figure, two biological replicates and several technical replicates were used.

Based on the findings that SDRRs play central roles in *C. crescentus* development and physiological adaptation, we set out to genetically characterize select SDRRs in this organism. Here, we present evidence that the SDRR MrrA is an important regulator of developmental processes such as motility and attachment and that it controls cell behavior through LovK. Importantly, MrrA is also a central component of the *C. crescentus* general stress response that directly impacts on PhyK and PhyR phosphorylation. Our results demonstrate that MrrA shuttles phosphoryl groups from a range of upstream kinases to both LovK and PhyK, which serve as histidine phosphotransferases. Based on these findings, we postulate that MrrA serves as a central phosphorylation hub that coordinates developmental processes with the general stress response in this organism.

## Results

### MrrA is a response regulator that controls development and the general stress response

To functionally characterize SDRRs in *C. crescentus*, in-frame deletions were generated in the respective genes (CC0630, CC2576, CC3015, and CC3286). These four SDRRs were chosen from a total of 20 SDRRs in *C. crescentus* based on the prediction that they are not involved in chemotaxis and had not been functionally characterized before (10, 15, 18, 23, 25). Whereas mutations of CC0630, CC2576 or CC3286 showed no apparent phenotype in the assays tested, a strain lacking CC3015 showed several behavioral and growth defects, based on which we renamed this protein MrrA for <u>m</u>ultifunctional <u>r</u>esponse <u>r</u>egulator <u>A</u>. When grown in a complex medium (PYE) the Δ*mrrA* strain showed wild type-like growth in exponential phase but entered stationary phase prematurely (Figure 1B). This suggested that the Δ*mrrA* mutant may lack the ability to cope with certain forms of stress associated with stationary phase. To better understand the mechanisms provoking this phenotype, we compared the proteomes of *C. crescentus* wild-type and Δ*mrrA* strains. Strong reductions in protein abundance in the *mrrA* mutant were observed for central components of the general stress response pathway, including PhyR, NepR and SigT, as well as for proteins that were previously identified as targets of SigT, including the histidine kinase LovK (Figure 1C, Table S1) (4, 25). Because the core components of the general stress response are subject to autoregulation (Figure 1A) (26, 28) these changes suggested that the Δ*mrrA* mutant failed to induce the general stress response under these growth conditions. This idea is consistent with the observation that the relative loss of fitness of a *mrrA* and a *sigT* mutant correlate for a range of different conditions (Figure S1A) (30). To verify an role of MrrA in the general stress response we made use of a *sigU*-*lacZ* reporter fusion, the activity of which strictly depends on the sigma factor SigT (25). In *C. crescentus* wild type, SigT was active in exponential phase and induced under osmotic stress or upon entry into stationary phase. In contrast, the Δ*mrrA* mutant showed no SigT activity irrespective of growth phase and stress applied (Figure 1D), arguing that MrrA is indispensable for the activity of SigT. In line with this notion, the Δ*mrrA* strain showed a 1000-fold reduction in survival as compared to wild type when challenged by oxidative stress, essentially phenocopying a *sigT* null mutant (Figure 1E).

The Δ*mrrA* mutant also displayed increased surface attachment and increased spreading on semisolid agar plates (Figure 1F), the latter of which requires an intact flagellar machinery and chemotaxis behavior. This indicated that MrrA, directly or indirectly, inhibits both motility/chemotaxis and holdfast-dependent attachment, two behaviors that are usually regulated inversely (31). A role for MrrA in holdfast production was further supported by the observation that the Δ*mrrA* mutant showed a strong increase in number and size of rosettes, characteristic holdfast-mediated *C. crescentus* cell aggregates (Figure S1B) (23, 25). Because both motility and attachment are regulated by the second messenger c-di-GMP (31), we determined c-di-GMP levels in the Δ*mrrA* strain but found no significant differences as compared to wild type (Figure S1C). Holdfast biogenesis is also regulated by the holdfast inhibitor protein HfiA, a component that directly binds to and inhibits HfsJ, an essential component of the adhesive polysaccharide export machinery (32). Moreover, LovK and LovR were shown to regulate *C. crescentus* surface attachment via the expression of HfiA (32). Epistasis experiments revealed that the effect of MrrA on attachment depended on LovK but not on LovR (Figure 1G). Likewise, *hfiA* expression was reduced in the Δ*mrrA* mutant, an effect that was dependent on LovK but not on LovR (Figure 1H). These results support the notion that MrrA acts upstream of LovK to control attachment via the modulation of *hfiA* expression.

In sum, these results revealed MrrA as a pleiotropic regulator affecting motility, attachment, growth and survival under stress conditions. In particular, our data indicated that MrrA is an essential component of the alpha-proteobacterial general stress response in *C. crescentus*. In the following we focused on unraveling the molecular mechanism by which MrrA impacts on the general stress response pathway.

### MrrA is a central phosphorylation hub for multiple histidine kinases

SDRRs function in phosphotransfer reactions or as allosteric regulators via protein-protein interaction. To identify MrrA interaction partners or phospho-donors we used yeast-two-hybrid screening and co-immunoprecipitation. These approaches identified several candidate proteins including the three histidine kinases CC2501, CC2554 and CC2874 (Table S1). *In vitro* phosphorylation assays with purified proteins failed to show phosphotransfer from CC2501 to MrrA (data not shown) but demonstrated rapid transfer from CC2554 and CC2874 to the conserved Asp53 residue of MrrA (Figures 2A, S2A). Moreover, when ATP was depleted upon addition of hexokinase and glucose (33), MrrA was efficiently dephosphorylated (Figure 2B). Although CC2874 harbors a C-terminal Rec domain, this part of the kinase is not required for autophosphorylation and phosphotransfer to MrrA (Figure S2A). These experiments demonstrated that CC2874 and CC2554 are cognate histidine kinases of MrrA and that both enzymes are able to dephosphorylate MrrA upon ATP depletion.

**Figure 2:**
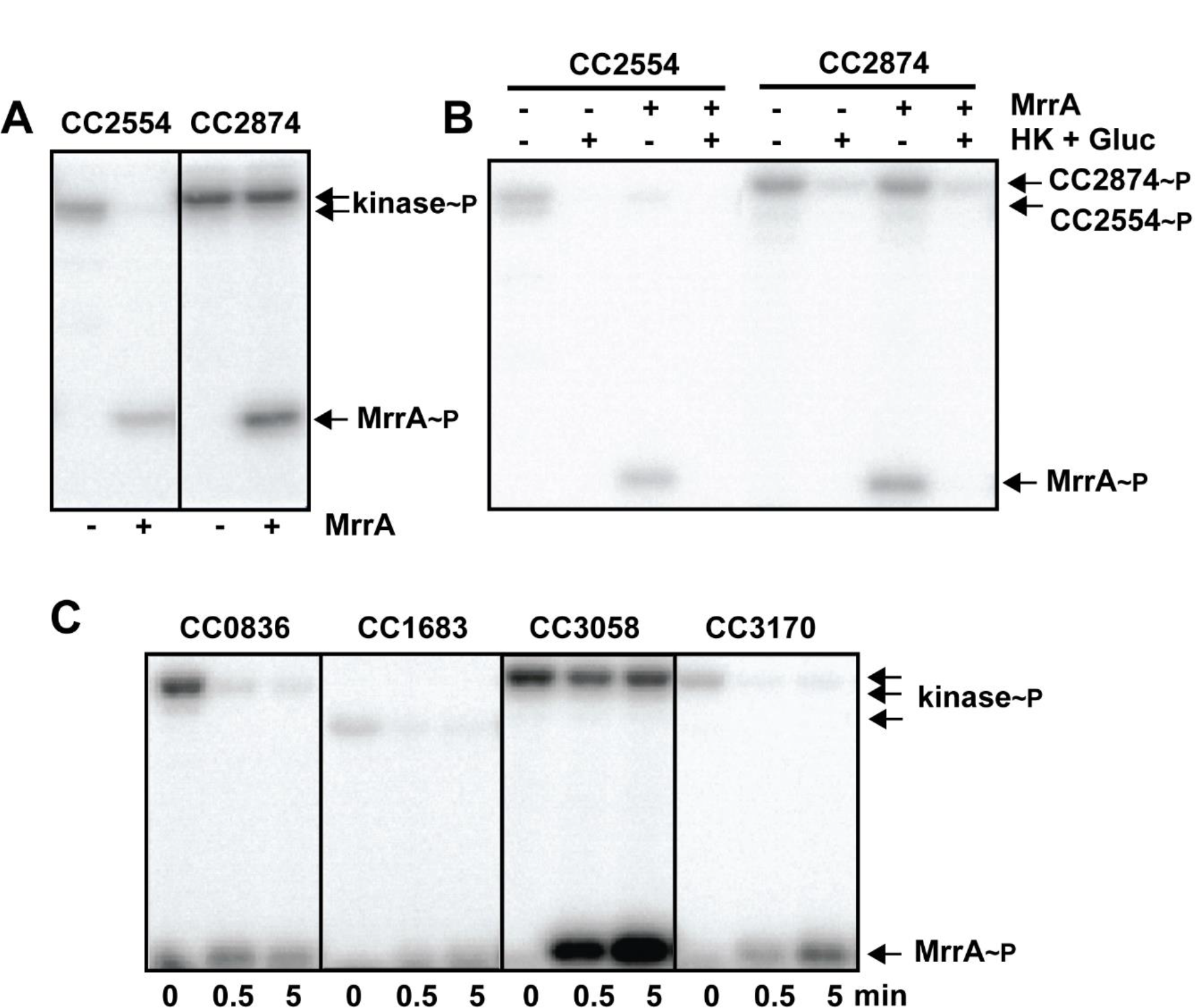
MrrA is phosphorylated by several upstream kinases. **(A)** Autophosphorylation of histidine kinases CC2554 (left) and CC2874 (right) and phosphotransfer to MrrA. Kinases (5 µM) and MrrA (10 µM) were mixed with 500 µM ATP and 2.5 µCi [γ^32^P]ATP (3,000 Ci mmol^-1^) as indicated. Reactions were carried out for 15 minutes at room temperature and analyzed by SDS-PAGE and autoradiography. The position of phosphorylated proteins on the gel is indicated on the right. **(B)** ATP depletion results in back-transfer of phosphate from MrrA to CC2554 (left) and CC2874 (right). Kinase reactions were incubated with and without MrrA as indicated in (A) for 15 minutes. ATP was depleted by adding hexokinase (1.5 units) and D-glucose (5 mM). The position of phosphorylated proteins on the gel is indicated on the right. **(C)** Several histidine kinases of the HWE subfamily transfer phosphate to MrrA. Purified kinases CC0836, CC1683, CC3058, CC3170 were mixed with ATP as indicated in (A) and auto-phosphorylation reactions were carried out for 30 min. MrrA was then added to the reactions and samples were taken at the time points indicated. The position of phosphorylated proteins on the gel is indicated on the right.

A recent study had implicated the SDRR SdrG in the general stress response of the alpha-proteobacterium *Sphingomonas melonis* Fr1 (34). Sequence comparison revealed that SdrG and MrrA harbor a conserved PFXFATG(G/Y) motif that distinguishes these proteins from prototypical response regulators (35). Moreover, SdrG and MrrA are best bidirectional hits in BLAST searches, indicating that SdrG and MrrA are orthologs. SdrG is phosphorylated by most members of the HisKA2 and HWE subfamilies of histidine kinases in *S. melonis* (34). Likewise, CC2554 is a member of the HWE subfamily, while PhyK and LovK, two proteins that have previously been implicated in the general stress response pathway in *C. crescentus*, belong to the HisKA2 subfamily (Figure S2B) (25, 29). This prompted us to test if other HisKA2/HWE kinases of *C. crescentus* are able to phosphorylate MrrA. LovK, PhyK and the remaining nine HWE/HisKA2 kinases of *C. crescentus* (CC0629, CC0836, CC1683, CC2909, CC3048, CC3058, CC3170, CC3198 and CC3569) (Figure S2B) were purified and analyzed for autophosphorylation and phosphotransfer to MrrA. Of the eleven kinases tested, five showed robust autophosphorylation under the conditions used. Four of the five kinases that were active *in vitro*, showed rapid phosphotransfer to MrrA (Figure 2C, Figure S2C). Notably, among the kinases lacking autokinase activity were also PhyK and LovK (see below).

Altogether, these results demonstrated that MrrA is phosphorylated by multiple HWE and HisKA2 histidine kinases. Kinase CC2874, a classical HisKA histidine kinase that does not belong to the HWE or HisKA2 subfamilies, also efficiently phosphorylated MrrA. These data established MrrA as a central phosphorylation hub that collects phosphate from members of different kinase families to control *C. crescentus* general stress response activity.

### MrrA controls the activation of the general stress response proteins PhyK and LovK

Previous studies demonstrated that the histidine kinase PhyK is essential for the general stress response in *C. crescentus* in vivo (25, 29). In contrast, LovK was proposed to be a negative regulator of the general stress response by promoting dephosphorylation of PhyR (25) (Figure 1A). However, biochemical evidence for the catalytic activity of PhyK as a genuine histidine kinase and for a role of LovK and PhyK in PhyR (de)phosphorylation is missing. Since the above genetic studies identified MrrA as an activator of the general stress response, we wondered whether MrrA could act as a direct activator of PhyK or LovK. *In vitro* phosphorylation experiments revealed that PhyK and LovK were readily phosphorylated in the presence of CC2874, MrrA and ATP, but not when incubated with ATP alone (Figure 3A-C). Next, we tested if LovK and PhyK could pass on phosphoryl groups to PhyR. Because it was previously shown that PhyR of *S. melonis* was efficiently phosphorylated by cognate histidine kinases only in the presence of the anti-sigma factor NepR, NepR was included in the phosphotransfer reactions containing PhyR (34). Although both PhyK and LovK were able to phosphorylate PhyR under these conditions (Figure 3A,B), phosphorylation by PhyK was rapid and efficient, while phosphorylation by LovK was slow and comparably weak (Figure 3D,E). These results are in line with the notion that PhyK, but not LovK, is the primary phospho-donor for PhyR (Figure 1A).

**Figure 3:**
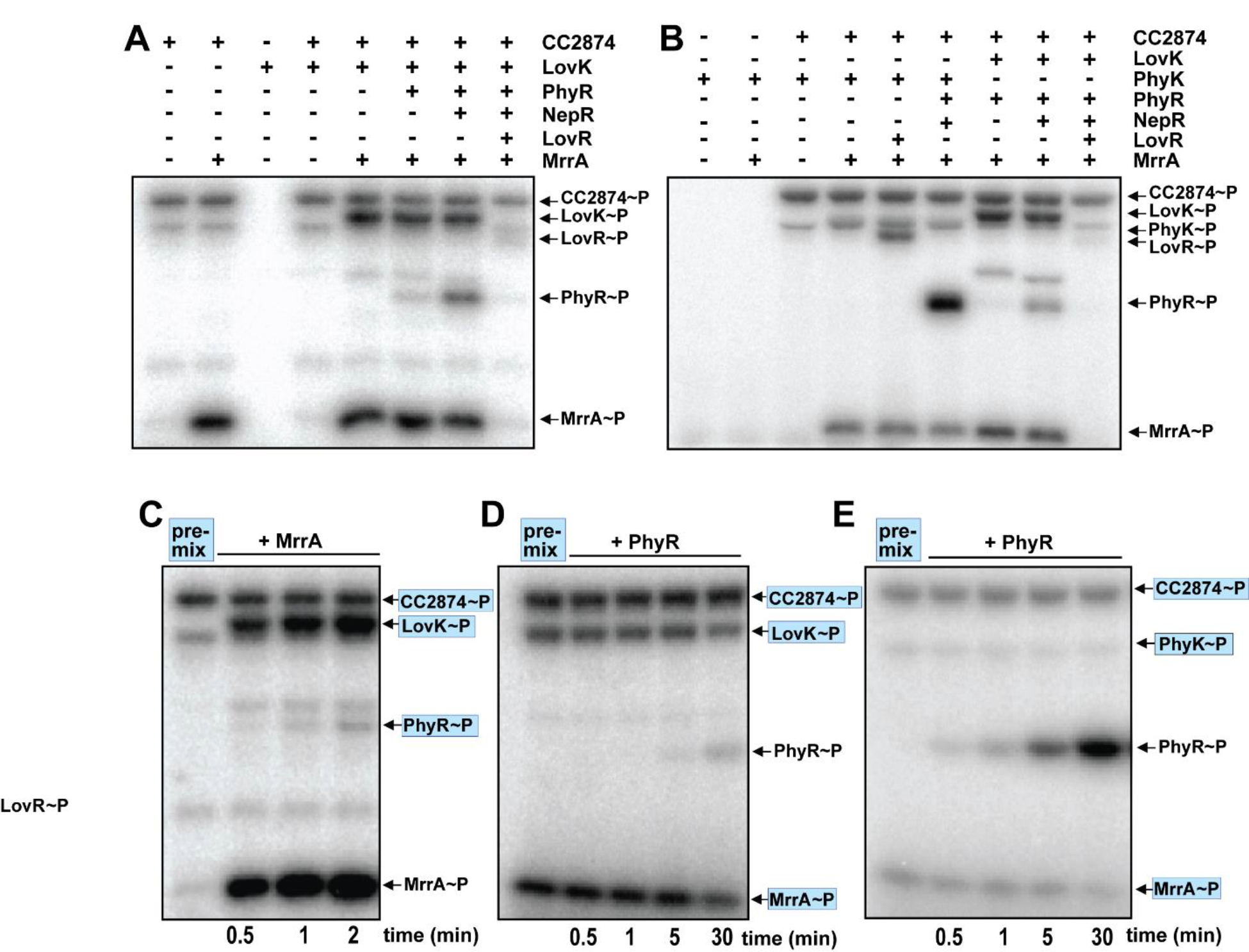
MrrA phosphorylates components of the general stress response. **(A)** MrrA transfers phosphate to LovK and PhyR. 500 µM ATP and 2.5 µCi [γ^32^P]ATP (3,000 Ci mmol^-1^) were mixed with the proteins indicated and reactions were carried out for 15 min at room temperature and analyzed by SDS-PAGE and autoradiography. The position of phosphorylated proteins on the gel is indicated on the right. Note that LovK does not display autokinase activity by itself but is readily phosphorylated if MrrA is present. **(B)** MrrA transfers phosphate to PhyK and PhyR. Phosphorylation reactions were assembled and run as in (A). The results obtained for PhyK were similar to the results obtained for LovK. Note that PhyK does not display autokinase activity but is readily phosphorylated if MrrA is present. **(C)** Phosphorylation of MrrA and LovK is rapid, while but phosphorylation of PhyR is slow. Phosphorylation reactions with radiolabeled ATP and purified CC2874, LovK, PhyR and NepR were pre-incubated for 30 minutes (pre-mix). Purified MrrA was then added to the reactions and samples were taken at the time points indicated. The position of phosphorylated proteins on the gel is indicated on the right. **(D)** PhyR phosphorylation through LovK is slow and inefficient. Phosphorylation reactions with radiolabeled ATP and purified CC2874, LovK, MrrA and NepR were pre-incubated for 30 minutes (pre-mix). Purified PhyR was then added to the reactions and samples were taken at the time points indicated. The position of phosphorylated proteins on the gel is indicated on the right. **(E)** PhyR phosphorylation through PhyK is rapid and efficient. Phosphorylation reactions with radiolabeled ATP and purified CC2874, PhyK, MrrA and NepR were pre-incubated for 30 minutes (pre-mix). PhyR was then added to the reactions and samples were taken at the time points indicated. The position of phosphorylated proteins on the gel is indicated on the right.

The data above suggested that phosphorylated MrrA acts as a direct activator of PhyK and LovK, explaining the strong stress response phenotype observed for the Δ*mrrA* strain *in vivo*. We reasoned that MrrA~P could either allosterically activate PhyK and LovK kinase activities (10) or serve as phospho-donor for PhyK and LovK (Figure 4A). The latter scenario would imply that PhyK and LovK do not serve as kinases but have adopted a role as histidine phosphotransfer proteins (HPt). In fact, DHp domains of histidine kinases and of HPt domains are structurally very similar (36–38). To distinguish between the two possibilities, we purified variants of PhyK harboring mutations in conserved residues of the G1 or G2 boxes of its catalytic (CA) domain that are essential for ATP binding. If PhyK is a *bona fide* kinase that is allosterically activated by MrrA~P, mutations in the ATP-binding pocket should abolish autophosphorylation (6). However, not only did both mutant proteins still accumulate radiolabel in a CC2874- and MrrA-dependent manner (Figure 4B), but they were also able to phosphorylate PhyR (Figure 4C). Together, this argued that PhyK serves as a phosphotransferase to shuttle phosphate from MrrA to PhyR. Similarly, LovK G1 and G2 mutant variants were phosphorylated indistinguishable from wild-type LovK in a reaction that required the kinase CC2874 and MrrA (Figure 4D).

**Figure 4:**
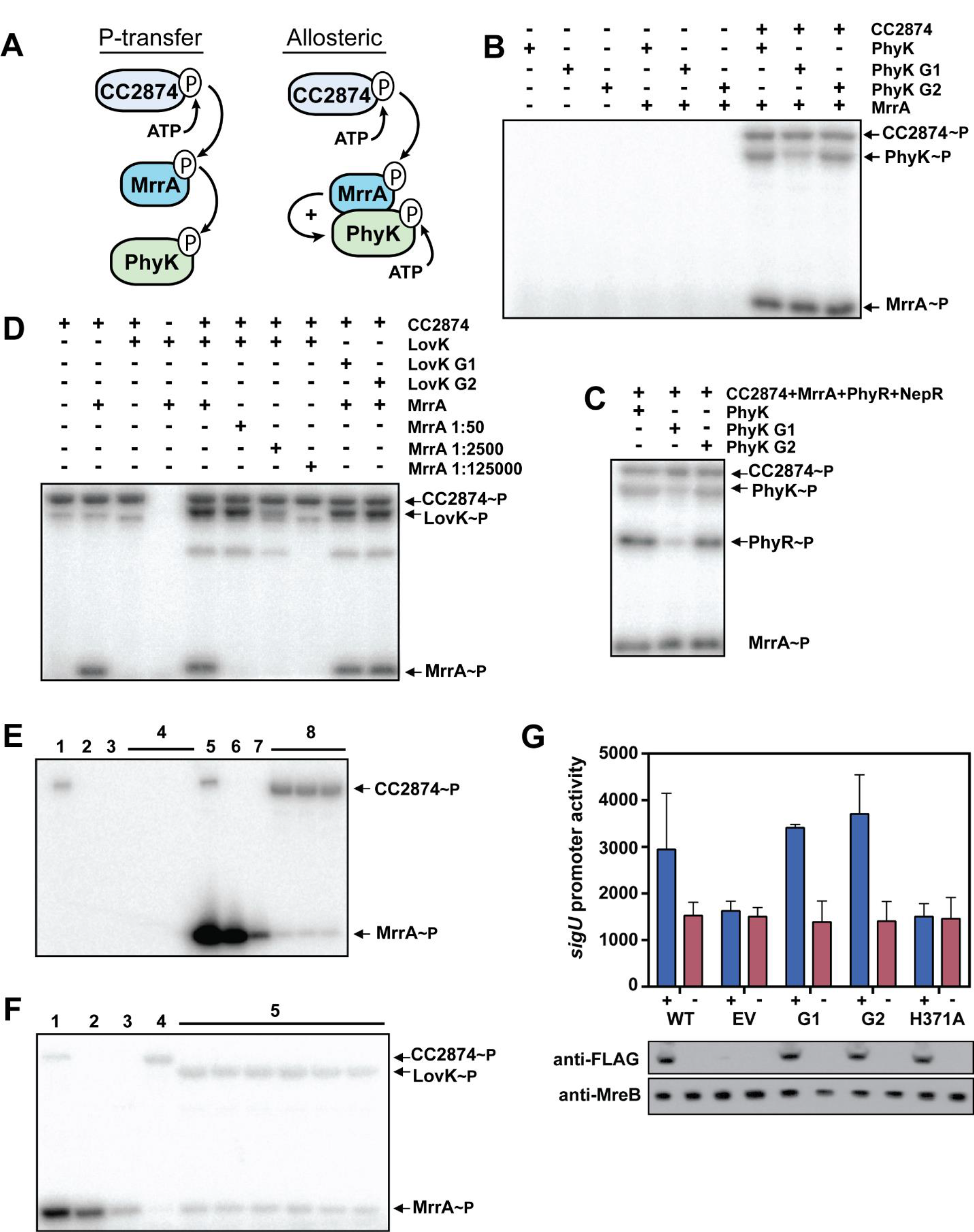
LovK and PhyK are phosphotransferases that are activated by MrrA~P. **(A)** Schematic representation of two possible modes of action of MrrA~P, phosphotransfer to and allosteric activation of PhyK and LovK. For reasons of simplicity only PhyK is shown. **(B)** PhyK phosphorylation does not require a conserved CA domain. Phosphorylation reactions with radiolabeled ATP, the kinase CC2874, MrrA, and different PhyK variants. Purified PhyK wild type or mutant variants harboring mutations in the G1 (G514A/G516A) or G2 (G526A) bo× of the ATP binding site were used as indicated. The position of phosphorylated proteins on the gel is indicated on the right. **(C)** LovK phosphorylation does not require a conserved CA domain. Phosphorylation reactions with radiolabeled ATP, kinase CC2874, MrrA, and different LovK variants. Purified LovK wild type or mutant variants harboring mutations in the G1 (G319A/G321A)) or G2 (G332A) bo× ATP binding site of the CA domain, were used as indicated. MrrA dilution factors are indicated in each lane. The position of phosphorylated proteins on the gel is indicated on the right. **(D)** Phosphotransfer to PhyR does not require a conserved CA domain. Phosphorylation reactions containing radiolabeled ATP, kinase CC2874, MrrA, PhyR, NepR and different PhyK variants are as in (C). The position of phosphorylated proteins on the gel is indicated on the right. **(E)** Preparation of purified MrrA~P. Kinase CC2874 alone or with MrrA were phosphorylated with radiolabeled ATP (lanes 1, 5) and CC2874 was subsequently removed using anti-MBP magnetic beads (lanes 2, 6). Next, ATP was hydrolyzed by treating mixtures with hexokinase and glucose (lanes 3, 7). CC2874 was added back to ATP-depleted samples and mixtures were incubated for 0.5, 1.0, and 5 minutes (lanes 4, 8). **(F)** Purified MrrA~P transfers phosphate to LovK. MrrA~P was prepared as in (E) (lane 1), CC2874 was removed (lane 2) and ATP was degraded (lane 3). Fresh CC2874 (lane 4) or LovK was added and phosphotransfer from MrrA~P was monitored after 10, 20 sec, 1, 2, 10 and 20 minutes (lanes 5). **(G)** ATP binding is not required for PhyK activity in the general stress response. Δ*lovK* strains harboring an empty vector (EV) or a plasmid expressing different *phyK* alleles from a cumate-inducible promoter were analyzed. Plasmid-driven variants of PhyK contained mutations in the G1 or G2 bo× of the ATP binding pocket (see above) or in the conserved phospho-acceptor His371. SigT-dependent *sigU* promoter activity (Miller units) was determined using a *lacZ* promoter fusion in strains grown in the presence (+) or absence (-) of cumate. PhyK variants harbored a C-terminal 3xFLAG-tags that allowed monitoring their expression by immunoblot analysis (lower panels). An immunoblot with anti-MreB antibodies is shown as control. Note that the *sigUp-lacZ* reporter fusion used in these experiments differed from the one used in experiments above (Fig. 1D) and shows higher basal activity (compare wild-type PhyK and empty vector control in Figure 4G).

To corroborate the idea that LovK and PhyK do not act as kinases but rather as HPt-like proteins, we established a procedure to purify radiolabeled MrrA~P, in order to directly follow phosphotransfer from MrrA to LovK or PhyK in the absence of ATP. To this end, MrrA was phosphorylated *in vitro* using MBP-tagged CC2874 and radiolabeled ATP. MBP-CC2874 was then removed from the reaction mix with anti-MBP magnetic beads and the remaining ATP depleted from the MrrA~P preparation using hexokinase and glucose. As a control, the same procedure was performed without MrrA. When CC2874 was added to this mock preparation, it failed to accumulate radiolabel, indicating that ATP was efficiently removed (Figure 4E). In contrast, CC2874 was readily phosphorylated when incubated with the preparation containing MrrA~P, indicating back-transfer from MrrA~P to CC2874 (Figure 4E). Similarly, when MrrA~P was incubated with LovK or PhyK, rapid accumulation of radiolabel on both proteins was observed (Figures 4F, S3A). No radiolabel accumulated on LovK or PhyK when incubated with cold, i.e. non-radiolabeled MrrA~P, even though radiolabeled ATP was present in the reaction mixture (Figure S3B). Together, these results strongly implied that MrrA acts as a shuttle to transfer phosphate from the kinase CC2874 to LovK and PhyK. These experiments also suggested that LovK and PhyK do not primarily act as kinases, but rather serve as phosphotransfer proteins to control the activity of PhyR and, ultimately, to control the activity of SigT.

To test the physiological relevance of our biochemical data, we sought to analyze general stress response activity of *C. crescentus* strains harboring mutations in the G1 and G2 boxes of PhyK. To this end, 3xFLAG-tagged mutant variants of PhyK were expressed *in trans* from a cumate-inducible promoter in a Δ*phyK* strain and SigT activity was monitored using a *sigU*-*lacZ* reporter. In line with our biochemical data, both PhyK mutants fully complement the Δ*phyK* phenotype (Figure 4G). In contrast, PhyK with a mutation of the conserved phospho-acceptor histidine (H371A) failed to rescue SigT activity. Thus, ATP binding and autokinase activity is not required for PhyK activity *in vivo*, strengthening the notion that it acts exclusively as a phosphotransfer protein to promote PhyR phosphorylation.

### LovR is a selective phosphate sink for MrrA, but not for PhyR

In the experiments described above, we established that MrrA samples information from multiple upstream kinases and, in response, shuttles phosphoryl groups to both LovK and PhyK. Whereas these observations are in good agreement with the genetic data demonstrating that MrrA and PhyK are part of the *C. crescentus* general stress response, they do not account for the described negative effect of LovK on the general stress response (25). Strains lacking LovK or LovR showed increase SigT activity and SigT-dependent survival under stress conditions; moreover, co-overexpression of *lovK* and *lovR*, but not of *lovK* or *lovR* alone, abolished SigT activity (25). To explain these genetic results and to rationalize how LovK and LovR downregulate SigT activity, it was proposed that LovK, together with LovR, could serve to promote PhyR dephosphorylation. Such a mechanism could be based on i) LovK serving as phosphatase of PhyR with LovR as the terminal phosphate sink (Figure 5, model 1); or on ii) phosphotransfer from PhyK to LovR with LovK acting as a LovR phosphatase (Figure 5, model 2) (25). Considering our findings that MrrA is positioned upstream of PhyK and LovK, we reasoned that the role of LovK and LovR may be to drain phosphate away from the PhyK-PhyR branch by re-routing phosphate flux via MrrA (Figure 5, model 3).

**Figure 5:**
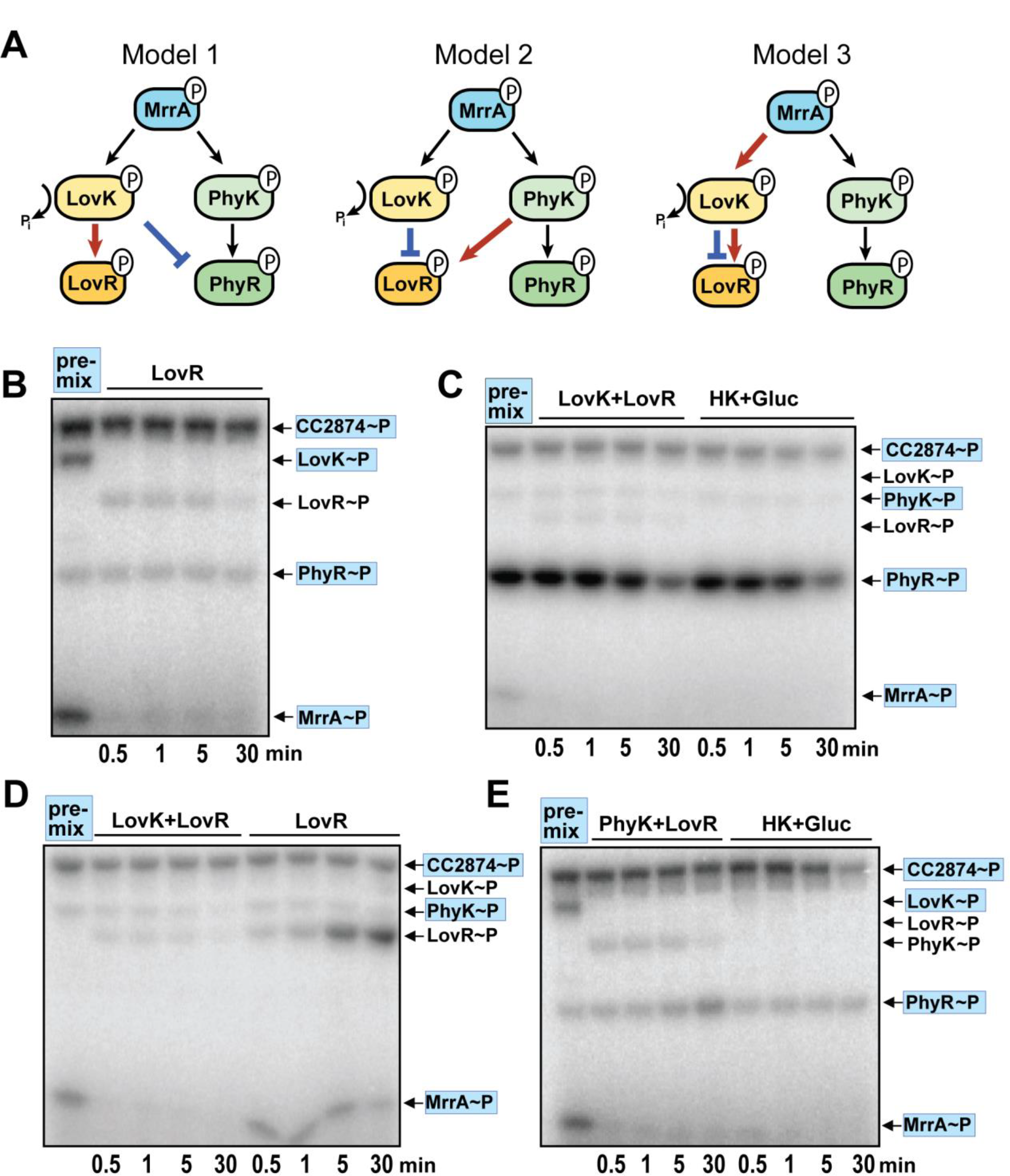
LovK and LovR rapidly dephosphorylate MrrA, but not PhyK or PhyR. **(A)** Models for the role of LovK and LovR in the regulation of the *C. crescentus* general stress response. PhyR activity could be reduced by LovK acting as phosphatase for PhyR~P with LovR as terminal phosphate sink (model 1). PhyK activity could be reduced by direct phosphotransfer to LovR with LovK acting as phosphatase for LovR~P (model 2). MrrA activity could be reduced by phosphotransfer to LovK and LovR. In this scenario LovR serves as terminal phospho-acceptor with LovK serving both as HPt and as phosphatase for LovR. **(B)** LovK is not a phosphatase for PhyR. Phosphorylation reactions with radiolabeled ATP and purified CC2874, LovK, PhyR, MrrA and NepR were incubated for 30 minutes (pre-mix). Phosphorylation levels of all proteins were monitored before and after addition of LovR at time points indicated. **(C)** PhyK does not transfer phosphate to LovR and LovK. Phosphorylation reactions with radiolabeled ATP and purified CC2874, MrrA, PhyR and NepR were incubated for 30 minutes (pre-mix). Phosphorylation levels of all proteins were monitored before and after addition of LovK and LovR or after addition of hexokinase and glucose at time points indicated. **(D)** Lov~P levels accumulate in the absence of LovK. Phosphorylation reactions with radiolabeled ATP and purified CC2874, MrrA and PhyK were incubated for 30 minutes (pre-mix). Phosphorylation levels of all proteins were monitored before and after addition of LovK and LovR or after addition of LovR alone at time points indicated. **(E)** PhyK does not transfer phosphate to LovR and LovK. Phosphorylation reactions with radiolabelled ATP and purified CC2874, MrrA, LovK, PhyR and NepR were incubated for 30 minutes (pre-mix). Phosphorylation levels of all proteins were monitored before and after the addition of PhyK and LovR or glucose/hexokinase at the time points indicated. **(D)** Phosphorylation reactions with radiolabelled ATP and purified CC2874, PhyK, MrrA, and NepR were incubated for 30 minutes (pre-mix). Phosphorylation levels of all proteins were monitored before and after the addition of LovK/LovR or after addition of hexokinase and glucose at time points indicated. The position of phosphorylated proteins on the gel is indicated on the right.

When PhyR was phosphorylated in the presence of CC2874, MrrA and LovK, addition of LovR resulted in the instant loss of LovK~P and MrrA~P, but not PhyR~P (Figure 5B). Notably, under these conditions only weak accumulation of LovR~P was observed, arguing that LovR~P is subject to rapid dephosphorylation. Thus, LovK does not serve as phosphatase for or phospho-acceptor of PhyR~P *in vitro*. To test if PhyR was dephosphorylated via PhyK and LovR, PhyR was phosphorylated in the presence of CC2874, MrrA and PhyK and LovK and LovR were added to the reaction mixture. This led to an instant loss of MrrA~P, but not of PhyR~P or PhyK~P (Figure 5C). When LovR alone was added to the pre-mix, MrrA~P was not depleted and instead LovR~P accumulated (Figure 5D). These observations support the idea that LovK and LovR function together to drain phosphoryl groups from MrrA, and that LovR~P does not undergo spontaneous dephosphorylation but that LovK acts as phosphatase for LovR~P. Similarly, when PhyR was phosphorylated in the presence of CC2874, MrrA and LovK, addition of PhyK and LovR led to a rapid loss of MrrA~P and LovK~P, but not of PhyR~P (Figure 5E). Finally, we tested if PhyR~P could be dephosphorylated upon depletion of ATP by hexokinase treatment and the resulting switching of CC2874 into phosphatase mode. Irrespective of how PhyR was phosphorylated, both PhyR and PhyK retained the radiolabel, while MrrA~P and LovK~P were rapidly dephosphorylated under these conditions (Figure 5C,E).

In conclusion, these results strongly argue against models 1 and 2 in Figure 5A but support model 3, in which the negative effect of LovK and LovR on the general stress response is a direct result of draining phosphate from MrrA. In contrast, cross-phosphorylation reactions between the LovKR and the PhyKR branch are unlikely. Importantly, rather than acting as a prototypical phosphatase of MrrA~P, LovK seems to act as phosphotransferase to shuttle phosphoryl groups from MrrA to LovR. In a final step, LovK then acts as a genuine phosphatase for LovR~P, a function that is essential to make LovKR an efficient phosphate sink.

## Discussion

Single-domain response regulators were previously shown to play important roles in *C. crescentus* cell cycle progression, development and behavior (15, 18, 23, 39). In the present study, we describe a novel single-domain response regulator in *C. crescentus*, MrrA, that is involved in a range of physiological processes, including growth, motility and attachment. Surprisingly, MrrA is also an integral and essential component of the general stress response in this organism. We propose a model, where MrrA is phosphorylated by several histidine kinases of different subclasses that are involved in the perception of diverse stress factors (25, 29, 34, 40). Once phosphorylated, MrrA shuttles phosphoryl groups to PhyK, which in turn phosphorylates PhyR, thereby inducing the partner switch triggering the general stress response (Figure 6). However, MrrA can also shuttle phosphate to LovK, a protein that, together with its cognate response regulator LovR, acts as a negative regulator of the general stress response. By draining phosphoryl groups from MrrA~P, LovK and LovR limit phosphorylation of PhyK and thus downregulate the general stress response. In addition, MrrA controls attachment by modulating the expression of *hfiA*, a process that is dependent on LovK, but the mechanistic details of this control are currently unclear. Altogether, this puts MrrA at the center of a complex signal transduction cascade that coordinates the *C. crescentus* stress response with behavioral adaptations.

**Figure 6:**
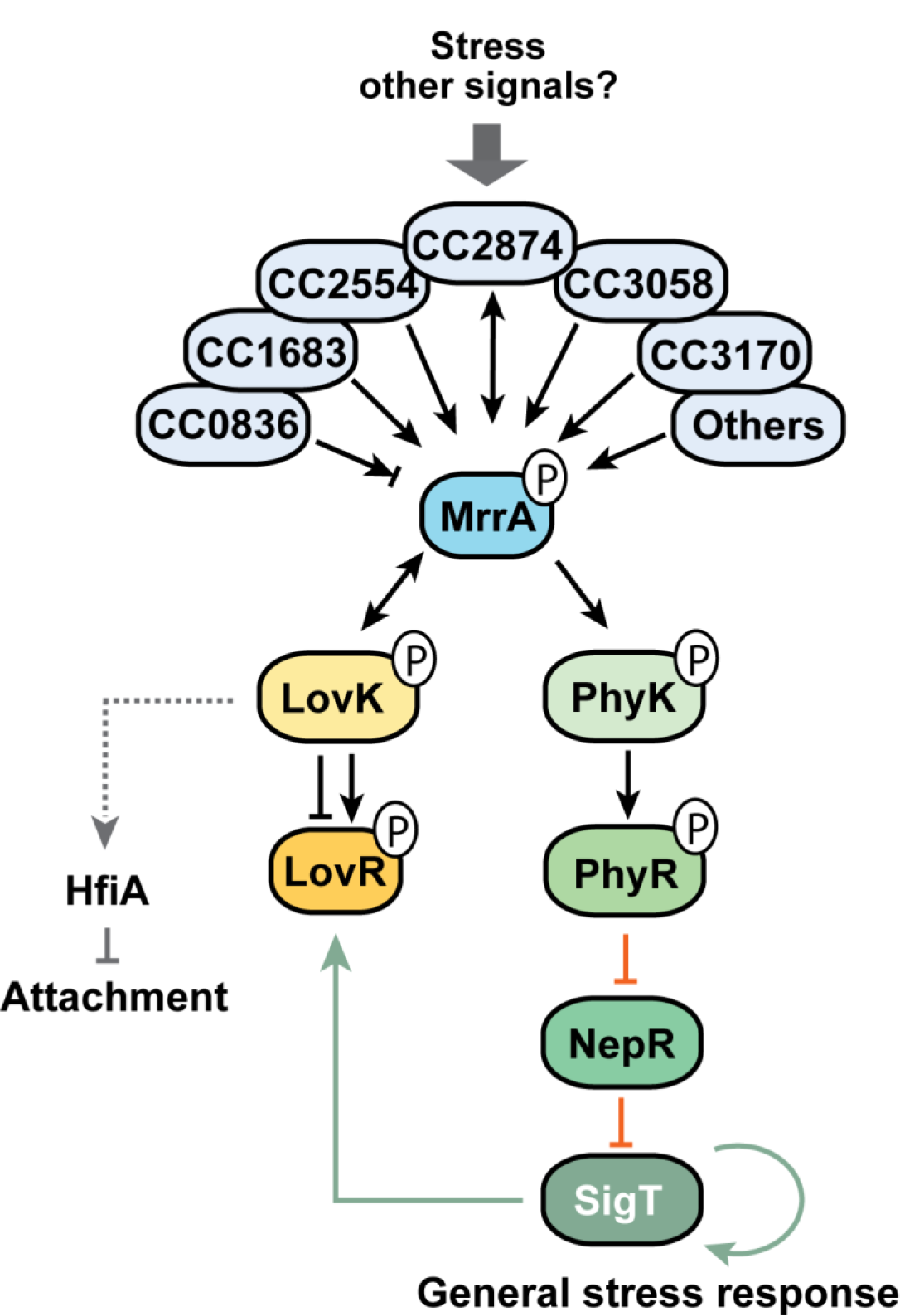
Model of MrrA function in the general stress response and developmental control of *C. crescentus*. We propose that members of different sub-families of histidine kinases sample different forms of stress and possibly other signals. Signaling converges through the phosphorylation the single domain response regulator MrrA. MrrA~P divergently distributes phosphate to the phosphotransfer proteins LovK and PhyK. Our data imply that PhyK~P is the principle phospho-donor for PhyR to activate the general stress response. LovK and LovR form a potent phospho-sink that involves LovK P-transfer to LovR, followed by rapid LovK-mediated removal of phosphoryl groups from LovR~P. Because *lovK* and *lovR* expression is SigT controlled, these proteins may constitute a feedback control mechanism that helps to adapt overshooting stress response. Alternatively, signals sensed by LovK may alter LovKR activity and by rerouting phosphate flu× from MrrA and fine-tune PhyR and ultimately SigT activity. Additional cellular processes like holdfast formation and surface attachment are integrated with the MrrA phosphorylation cascade through LovK. Black, green and red arrows indicate transcriptional regulation, phosphorylation and inhibitory protein-protein interactions, respectively. Grey arrows indicate regulatory links for which the mechanistic details are unknown.

### Multiple upstream histidine kinases phosphorylate MrrA

The observation that multiple histidine kinases of the HWE and HisKA2 subclasses phosphorylate MrrA, suggests that multiple signals are integrated at the level of MrrA phosphorylation. This notion seems reasonable since the general stress response is thought to respond to and protect from several unrelated stresses. In fact, in the related alpha-proteobacterium *S. melonis*, most HWE and HisKA2 kinases converge on PhyR phosphorylation with different kinases sensing different stresses. Interestingly, deletion of these kinases does not fully abrogate general stress response activity, suggesting that kinases outside of the HWE and HisKA2 subclasses also contribute to stress response activation (34). This is in agreement with our observation that MrrA is efficiently phosphorylated by the kinase CC2874, which belongs to the HisKA subfamily. We speculate that the number of kinases that can phosphorylate MrrA is even larger, since several HWE/HisKA2 kinases failed to auto-phosphorylate *in vitro* and because HisKA kinases were not systematically tested for phosphotransfer to MrrA.

The molecular cues that are sensed by the histidine kinases upstream of MrrA are currently unknown. Most kinases harbor classical sensing domains such as PAS and GAF domains, suggesting that they can sense stresses directly (Figure S2D). Although few kinases have been assigned specific sensory functions in the general stress response of alpha-proteobacteria, LOV domains sensing blue-light have been linked to this pathway in several organisms (25, 26, 28, 34, 41–45). LovK harbors a LOV domain and was previously shown to respond to blue light *in vitro* and to mediate attachment dependent on this stimulus (23). It is possible that phosphate flux through MrrA is modulated by blue light. In addition, the LOV domain of LovK was proposed to respond to redox conditions (25). This is based on the findings that a strain lacking PhyK and LovR but overproducing LovK still responds to oxidative stress. Although we cannot rule out this possibility, our data offer an alternative explanation for this observation, namely that an upstream kinase of MrrA responds to redox conditions and that LovK in this specific genetic context simply serves as an HPt shuttling phosphoryl groups to PhyR. A future challenge will be to assign sensory functions to individual kinases involved in the general stress response.

### PhyK is a phospho-transfer protein

Previous studies from independent groups have identified PhyK as a central component of the general stress response. It was assumed that PhyK acts as histidine kinase, which, in response to specific stimuli, phosphorylates the PhyR response regulator (25, 29). This assumption seemed plausible since PhyK has a conserved CA domain and harbors a putative periplasmic sensing domain. However, PhyK lacks autophosphorylation activity but rather functions as an HPt, accepting phosphate directly from MrrA~P. Moreover, conserved residues for ATP binding were dispensable for PhyK function *in vivo*. Finally, a Δ*mrrA* mutant quantitatively phenocopies a Δ*sigT* or Δ*phyK* strain in terms of its defect in general stress response activation, suggesting that in a Δ*mrrA* strain PhyK lacks residual kinase activity. It is currently unclear why the CA domain of PhyK (and LovK) is so well conserved despite its apparent lack of enzymatic activity. It is possible that the CA domain plays an allosteric role in signal transduction, contributes to structural stability and integrity or serves scaffolding functions. Of note, the *S. melonis* ortholog of PhyK, PhyP, harbors a degenerate CA domain and was proposed to act as a phosphatase of PhyR (46, 47). If PhyP also shuttles phosphoryl groups between PhyR and the MrrA orthologs, SdrG, in this organism is unknown. We also cannot rule out that PhyK has kinase activity under specific conditions, where it is able to directly respond to particular stresses. A previous study identified a cysteine residue in the periplasmic domain of PhyK that, when mutated, abolished PhyK function, suggesting that it is involved in stress sensing (29). If so, PhyK would be able to integrate multiple signals through its periplasmic sensor domain and through phosphorylation by MrrA~P. In fact, an earlier study reported autophosphorylation of LovK in vitro (23), although with our LovK expression construct and under the experimental conditions employed we do not observe a significant degree of LovK autophosphorylation. Importantly, our in vivo experiments clearly demonstrate that autokinase activity of PhyK is dispensable for PhyK function, strongly arguing that its essential function in the general stress response is that of a histidine phosphotransferase.

### Role of LovK and LovR as inhibitors of the general stress response

Recently, LovK and LovR were described as negative regulators of the general stress response (25). Different models were proposed for LovKR control that postulated dephosphorylation of PhyR as the mechanism to shut down the general stress response, predicting direct crosstalk between the PhyKR and LovKR branches (25). While our data confirmed that LovR can rapidly deplete phosphate from LovK, we did not observe dephosphorylation of PhyR~P or PhyK~P *in vitro*. Thus, the LovKR and PhyKR branches do not seem to crosstalk directly. Rather, LovKR restricts phosphate flow towards PhyKR by draining phosphoryl groups from the shared upstream component MrrA. Because NepR was always present during *in vitro* reactions, we cannot exclude that PhyR alone can be dephosphorylated by LovK or PhyK. However, this seems unlikely given the recent observation that NepR binding and PhyR phosphorylation are cooperative (48, 49). Since the expression of *lovKR* is positively controlled by SigT, LovKR likely constitute a negative feedback loop that serves to dampen the stress response (Figure 6). Our findings that LovK and LovR do not directly lead to PhyK or PhyR dephosphorylation imply that the dampening effect on SigT activity is provided by diverting phosphoryl groups away from MrrA and thus restricting future phosphorylation of PhyR. In combination with the stochiometric upregulation of the general stress response core components PhyR, NepR and SigT, this would ultimately result in the accumulation of unphosphorylated PhyR and the shut-down of the response (26, 28).

Our data show that MrrA~P can be dephosphorylated by at least two of its upstream kinases, CC2554 and CC2874, upon depletion of ATP (Figure 2B). Moroever, CC2874 directly accepts phosphoryl groups from MrrA~P (Figure 4E). We speculate that upstream components of MrrA are generally able to switch into phosphatase mode and shut down the response, possibly upon cessation of their respective input signals. Thus, the extent of MrrA phosphorylation may simply be dictated by mass action, i.e. free phosphate flow between up- and downstream kinases and MrrA, which in turn depends on whether kinases are in kinase or phosphatase mode. We have not quantitatively assessed MrrA auto-dephosphorylation in this study. However, based on the fact that we could easily isolate stable MrrA~P *in vitro* (Figure 4E), we propose that this process plays a minor role in phosphorelay dynamics. This is in good agreement with the presence of conserved Asn and Tyr residues at the D+2 and T+2 positions (50).

### Conserved and divergent roles of MrrA in alpha-proteobacteria

In this study, we show that MrrA is an essential and central component of the general stress response in *C. crescentus*. In contrast to prototypical response regulators, MrrA lacks one of the residues involved in the Y-T coupling mechanism required for intramolecular signal transduction of Rec domains. Instead, it harbors the recently described FATGUY motif (35, 51, 52). Three other FATGUY response regulators are described, SdrG of *S. melonis*, Mext_0407 of *Methylobacterium extorquens* and Sma0114 of *Sinorhizobium meliloti* (34, 53, 54). While the first two proteins are involved in the general stress response of these organisms, the latter is involved in succinate-mediated catabolite repression and polyhydroxybutyrate (PHB) production; whether or not Sma0114 also plays a role in the general stress response has not been tested. Hence, it seems reasonable to propose that MrrA and other members of this sub-family of response regulators play a conserved role in the general stress response of alpha-proteobacteria.

The degree to which MrrA orthologs contribute to general stress response activity seems to differ between organisms. In *S. melonis* and *M. extorquens*, deletion of the *mrrA* orthologs *sdrG* and Mext_0407 reduce, but do not completely abolish, general stress response activity (34, 53). In *S. melonis*, overexpression of most HWE/HisKA2 kinases leads to the induction of the general stress response. However, only a subset of these kinases require SdrG for this induction (34). Interestingly, in *S. melonis*, most HWE/HisKA2 kinases phosphorylate both PhyR and SdrG *in vitro*, arguing that specificity determinants of the receiver domains of PhyR and SdrG are similar (34, 35). These observations argue that the regulatory wiring of FATGUY response regulators to up- and downstream histidine kinases is plastic, similar to the plasticity and modularity of the sensory capacities of general stress response kinases themselves, likely reflecting species-specific niche adaptation (26, 28). In addition, FATGUY response regulators seem highly promiscuous with respect to their cognate kinases. This is in contrast to the vast majority of two-component systems that are thought to have evolved towards insulation (55, 56). Because known specificity determinants for histidine kinase-response regulator interaction were only identified for the HisKA subfamily and their prototypical response regulators employing Y-T coupling (57, 58), the molecular details of this promiscuity remain unknown. However, FATGUY response regulators like MrrA may have evolved as central phosphorylation hubs to integrate the phosphorylation status of multiple two-component systems, thereby coordinate the general stress response with cellular behavior and development. Pulldown experiments with MrrA (Table S1A) also identified the diguanylate cyclase DgcB (59, 60) and ChpT, a histidine phosphotransferase that plays a central role in cell cycle progression (18). It is possible that MrrA also intersects with c-di-GMP signaling and cell cycle control.

Networks combining multiple inputs with multiple output processes through a central “knot” are termed “bow tie”. Bow-tie architectures in metabolism or signal transduction were proposed to facilitate independent evolution of the input and output functions without affecting the regulatory core (61) and also compress cellular input information (62). One of the primary challenges for bacteria expanding their ecological niches is to quickly adapt to a plethora of novel stresses encountered at new sites and to effectively link this information with existing or emerging processes of stress response and behavior. We propose that the bow-tie architecture of phosphorylation network controlling stress response and behavior in *C. crescentus* ultimately facilitates niche adaptation.

## Materials and Methods

### Oligonucleotides, plasmids, strains and media

Oligonucleotides, plasmids and bacterial strains are listed in Table S2. *C. crescentus* was grown in PYE or M2G medium at 30°C (63). *E. coli* DH5α was used as host for cloning and grown in LB at 37°C. When required, the growth media were supplemented with antibiotics at the following concentrations (liquid / solid medium): 5/50 µg ml^-1^ of ampicillin, 5/20 µg ml^-1^ of kanamycin, 2.5/5 µg ml^-1^ of tetracycline, 1/2 µg ml^-1^ of chloramphenicol, 15/20 µg ml^-1^ of nalidixic acid (*C. crescentus*) and 50/100 µg ml^-1^ of ampicillin, 30/50 µg ml^-1^ of kanamycin, 12.5/12.5 µg ml^-1^ of tetracycline, 20/30 µg ml^-1^ of chloramphenicol, 15/30 µg ml^-1^ of nalidixic acid (*E. coli*).

### Growth experiments

Independent *C. crescentus* cultures were diluted to an OD_660_ of 0.05 in PYE medium. Three technical replicates (165 µl) of each culture were inoculated in 96-well plates and growth was monitored at 660 nm every 15 min in a Synergy H4 hybrid reader (BioTek) using Gen5 2.00 software (BioTek) at 30°C under shaking conditions (medium speed, continuous shaking).

### Attachment and motility assays

Surface attachment of *C. crescentus* was determined as described (64). Motility assays were carried out as described previously (15).

### Hydrogen peroxide stress assays

Stress assay was adapted from (65). Cells were grown overnight in M2X. Overnight cultures were diluted back to OD_660_ of 0.05. Cells were grown for 5 hours and again diluted back to OD_660_ of 0.05. Cultures were split and one culture was exposed to 0.2 mM of H_2_O_2_ for 1 hour. H_2_O_2_ (fresh bottle) was diluted back from a 10 mM solution. After the stress treatment cells were serially diluted 1:10 in M2X and spotted on PYE agar plates.

### Determination of c-di-GMP concentrations

C-di-GMP extraction and quantification was carried out as described previously (60).

### β-galactosidase assays

Independent *C. crescentus* cultures were grown in PYE to an OD_660_ of 0.3 (except for stationary phase cultures, which were taken directly from an ON culture). 2 ml culture was pelleted and resuspended in 2 ml fresh Z-buffer (0.06 M Na_2_HPO_4_, 0.04 M NaH_2_PO_4_, 0.01 M KCl, 0.001 M MgSO_4_, 0.3% β-mercaptoethanol). 1 ml was mixed with 100 µl of 0.1% SDS and 20 µl chloroform by vortexing for 10 sec and was incubated for 15-30 min. Three replicates of 200 µl each were transferred to a 96-well plate, 25 µl of fresh ONPG (β-D-galactopyranosid, 4 mg/ml stock) were added and β-galactosidase activity was measured in an EL800 plate reader (both Bio-Tek Instruments) over time and the maximum slope was plotted as increase of OD_405_ corrected for OD_660_ and volume. Alternatively, β-gal measurements were performed as described (66) using pAK504 and pAK505 *lacZ* reporter plasmids, either with exponentially growing cultures (*hfiA-lacZ* reporter fusion) or directly on overnight cultures (*sigUp-lacZ* reporter fusion) grown in PYE. Where appropriate, cumate was included in overnight cultures at a concentration of 100 µg/ml to induce gene expression from promoter P_Q5_ (46). pAK504 was constructed by ligation of a blunted *Aat*II/*Pci*I-fragment derived from pUT18 carrying *bla* and the ColE1 *oriV* into the *Xma*I site of pAK501 (46). To construct plasmid pAK502, part of *lacZ* was PCR-amplified from pAK501 using primers 9060/9061, the product was digested with *Kpn*I/*Dra*III and cloned into pAK501 digested with the same enzymes. The resulting plasmid encodes *lacZ* without a ribosome binding site and start codon and allows the construction of translational *lacZ* fusions. pAK505 was derived from pAK502 by subcloning a *Sac*II/*Eco*RI-fragment carrying *bla* and the ColE1 *oriV* from pAK504 in between the same sites of pAK502. Inserts were cloned in pAK504 and pAK505 using *Kpn*I and *Xba*I restriction sites and primers described in Table S2.

### Proteome analysis

Independent cultures of *C. crescentus* were grown to an OD_660_ of 0.3 and 10 ml of cells were pelleted and dissolved in 200 µl cold lysis buffer (8 M Urea, 0.1 M ammonium bicarbonate, 0.1% RapiGest). Cells were lyzed by ultrasonication (Vial Tweeter, Hielscher) (2× 10 sec, amplitude 100, cycle 0.5) and shaking in a Thermomixer C (Eppendorf, 5 min, 1400 rpm, RT). After centrifugation (30 min, 4°C, max speed), the supernatant containing the solubilized proteins was transferred to a fresh tube and the protein concentration was measured using a standard Bradford assay (Bio-Rad) and adjusted to a final concentration of 1 mg/ml. To reduce and alkylate disulfide bonds, 1 µl TCEP (tris(2-carboxyethyl)phosphine, 0.2 M stock in 0.1 M Tris pH 8.5) was added to 40 µl protein extract (37°C, 1h, 1000 rpm). After the samples were cooled down, 1 µl fresh iodacetamide solution (0.4 M stock in HPLC water) was added and incubated in the dark (25°C, 30 min, 500 rpm). Finally, 1 µl N-acetyl-cysteine solution (0.5 M stock in 0.1 M Tris pH 8.5) was added, vortexed and incubated (RT, 500 rpm, 10 min). For the proteolysis Lys-C (0.2 µg/µl stock, Wako) was added to a final enzyme/protein ratio of 1:100 (37°C, 4 h, 550 rpm). The sample was diluted 1:5 (v/v) to a final urea concentration below 2 M using fresh 0.1 M ABC buffer (ammonium bicarbonate in HPLC water). Porcine trypsin (0.4 µg/µl stock, Promega) was added to a final trypsin/protein ratio of 1:50 (37°C, ON, 550 rpm). Post digestion, TFA (trifluoroacetic acid, 5% stock in HPLC water) was used to decrease the pH below 2. For the solid phase extraction C18-microspin columns (Harvard Apparatus) were conditioned with 150 µl acetonitrile (2400 rpm, 30 sec) and equilibrated twice with 150 µl TFA (0.1% stock in HPLC water, 2400 rpm, 30 sec). The sample was transferred twice through the column (2000 rpm, 2 min) before the column was washed 5x with 150 µl wash buffer (5% acetonitrile, 95% HPLC water and 0.1% TFA) (2400 rpm, 30 sec). The peptides were eluted twice with 150 µl elution buffer (50% acetonitrile, 50% HPLC water and 0.1% TFA) and concentrated under vacuum to dryness using a table top concentrator (Eppendorf). The peptides were dissolved to a final concentration of 0.5 µg/µl in LS-MS/MS buffer (0.15% formic acid, 2% acetonitrile, HPLC water) using 20 pulses ultra-sonication (Vial Tweeter, Hielscher) (amplitude 100, cycle 0.5) and shaking in a Thermocycler (37°C, 5 min, 1.400 rpm) (Eppendorf).

### Co-immunoprecipitation analysis

Independent cultures of *C. crescentus* strains UJ5511 and UJ6643 were grown in PYE to an OD_660_ of 0.3. Cells were pelleted, washed twice in 50 ml 20 mM Tris pH 8.0, 100 mM NaCl, and resuspended in 10 ml Bug Buster (Novagen) supplemented with 1 µl complete mini protease inhibitor (Roche), 200 µg/ml lysozyme and benzonase (0.5 µl/ml). After incubation at room temperature (20 min, gentle shaking), cell debris was removed by centrifugation (10.000xg, 15 min, 4°C). 150 µl Protino® Ni-NTA Agarose (Macherey-Nagel) was washed 3x in 500 µl Bug Buster (1.000xg, 1 min, 4°C), and incubated with the cleared lysate (ON, 4°C, 10 rpm on a rotary wheel). The beads were transferred to a Biospin column (Bio-Rad) and washed 4x with 700 µl HNN-lysis buffer (50 mM HEPES pH 7.5, 150 mM NaCl, 50 mM NaF, 5 mM EDTA) with 0.5% IGEPAL CA-630 (Sigma-Aldrich), before being washed 4× with HNN-lysis buffer without detergent. The protein extract was eluted using 3x 150 µl 0.2 M glycine (in HPLC water, pH 2.5). The eluate was neutralized with 150 µl ABC-buffer (ammonium bicarbonate, 1M stock in HPLC grade water). Urea (8 M stock in 100 mM ABC-buffer) was added to a final concentration of 1.6 M and the sample was vortexed before reducing and alkylating disulfide bonds as follows. 1 µl TCEP (tris(2-carboxyethyl)phosphine, 0.2 M in 100 mM ABC-buffer) was added per 40 µl protein extract (37°C, 30 min, 1.000 xg). After cooling down, 1 µl fresh iodacetamide (0.4 M stock in HPLC water) was added per 40 µl protein extract and incubated in the dark (25°C, 30 min, 500 rpm). Finally, 1 µl N-acetyl-cysteine solution (0.5 M in 0.1 M ABC-buffer) was added per 40 µl sample, vortexed and incubated (25°C, 10 min, 500 rpm). For proteolysis 1 µg porcine trypsin (0.4 µg/µl stock, Promega) was added (ON, 37°C, 500 rpm). For peptide purification 150 µl TFA (trifluoroacetic acid, 5% stock in HPLC water) was added to decrease the pH below 3. C18-microspin columns (Thermo Scientific) were conditioned twice with 150 µl acetonitrile (1.600 rpm, 30 sec) and equilibrated 3x with 150 µl 0.1% TFA (2.400 rpm, 30 sec). The samples were loaded and the flow-through was collected in a fresh tube (1.800 rpm, 2 min). The flow-through was reloaded and centrifuged again (1.800 rpm, 2 min). A mixture of 5% acetonitrile, 95% HPLC water (v/v) and 0.1% TFA was used to wash the columns 3x with 150 µl volume (2.400 rpm, 30 sec). Bonded peptides were eluted into a new tube using 3x 100 µl elution buffer (50% acetonitrile, 50% HPLC water (v/v) and 0.1% TFA) (1600 rpm, 30 sec). A speed vac (Eppendorf) was used to concentrate the eluted peptide mixture to dryness. The peptides were dissolved in 50 µl LC-buffer A (0.15% formic acid, 2% acetonitrile) using 20 pulses ultra-sonication (Vial Tweeter, Hielscher) (amplitude 100, cycle 0.5) and shaking (25°C, 5 min, 1.400 rpm).

### Yeast two-hybrid screening

*S. cerevisiae* PJ69-4A (UJ5292) (67) was transformed with pIDJ041 (UJ6743). Single colonies of *S. cerevisiae* PJ69-4A containing the bait-plasmid pIDJ041 were used for library-scale transformation with a *C. crescentus* library (68). 2.2×10^6^ transformants were screened on plates lacking histidine (SC-Trp-Leu-His + 5 mM 3’AT) and single colonies were used to isolate prey plasmids for sequencing.

### Protein purification

*E. coli* BL21 containing plasmids of interest were grown at 30°C in Lysogeny Broth (LB) medium and induced with 1 mM isopropyl β-D-1-thiogalactopyranoside (IPTG) for protein overproduction at OD_600_ = 0.6. Cells were harvested 2 h after induction (5.000 rpm, 15 min, 4°C) and stored at -80°C. After resuspension in lysis buffer (1× PBS, 10 µg/ml DNase, 1 complete mini protease inhibitor tablet (Roche)), cells were disrupted by French pressing and the supernatant containing the protein of interest was separated from the cell lysate by centrifugation (11.000 rcf, 1 h, 4°C). The supernatant was incubated with 1 ml Protino® Ni-NTA Agarose (Macherey-Nagel) (11 rpm, 1 h, 4°C) and was washed with 2x PBS, 500 mM NaCl, 10 mM imidazole (pH 8.0), 1 mM DTT. The protein was eluted with 1 × PBS, 500 mM NaCl, 250 mM imidazole (pH 8.0), 1 mM DTT and dialyzed in Spectra/POR® membranes (Spectrum Laboratories) using 10 mM HEPES-KOH, pH 8.0, 50 mM KCl, 10% glycerol, 0.1 mM EDTA, pH 8.0, 5 mM β-mercaptoethanol, 5 mM MgCl_2_.

### *In vitro* phosphorylation

Kinase and phosphatase assays were adapted from (33). Unless otherwise stated, protein concentrations of 5 µM were used. Reactions were incubated in dialysis buffer in the presence of 500 µM ATP and 2.5 µCi [γ^32^P]ATP (3,000 Ci mmol^-1^, Hartmann Analytic) at room temperature. Additional proteins were added and reactions stopped by the addition of SDS sample buffer at indicated time points. Reactions were stored on ice or loaded on 12% SDS gels. Wet gels were exposed to phosphor screens (0.5-1.5 h) before being scanned using a Typhoon FLA7000 imaging system (GE Healthcare). In experiments assessing phosphatase activity, ATP was depleted by the addition of 1.5 units hexokinase (Roche) and 5mM D-glucose 15 minutes after phosphorylation. For the purification of MrrA~P the following conditions were used. CC2874 (0.2 µM) and MrrA (100 µM) were pre-phosphorylated for 1 hour. 25 µl of α-MBP magnetic beads New England BioLabs, E8037S) were added and incubated for 1 hour. Beads were then concentrated using a magnet. Hexokinase and glucose was added to the supernatant as described above and incubated for 10 min to deplete remaining ATP.

## Acknowledgements

We thank Fabienne Hamburger for help with cloning, Mohit Kumar for construction of strain UJ10487, Alexander Schmidt (Proteomics Core Facility, Biozentrum, University of Basel) for assistance with proteomics and Volkhard Kaever (Research Core Unit Metabolomics and Institute of Pharmacology, Hannover Medical School, Hannover, Germany) for c-di-GMP quantifications. This work was supported by the Swiss National Science Foundation (SNF) grant 310030B_147090 to U.J. and by an ERC Advanced Research Grant to U.J.

---

**Figure S1:**
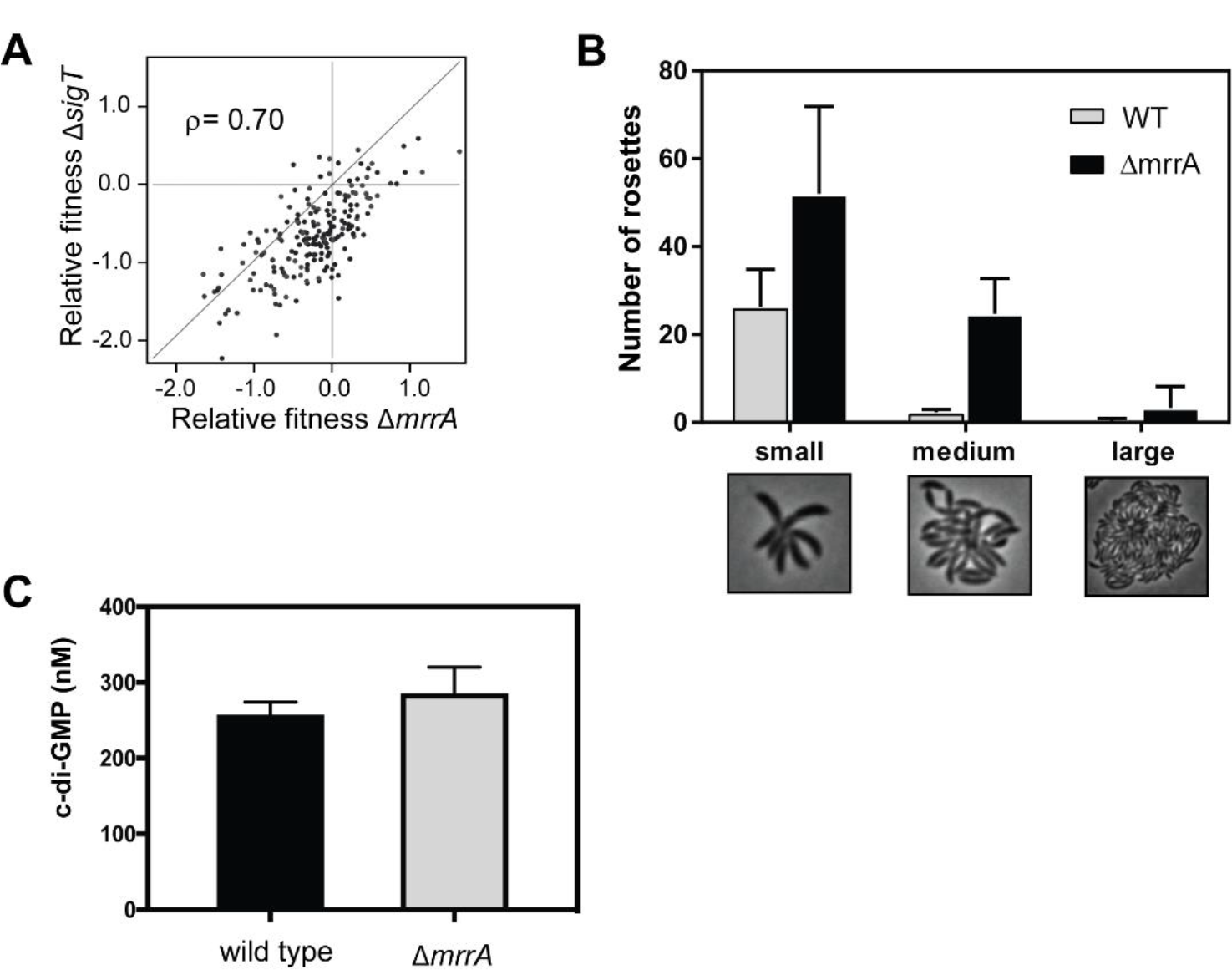
**(A)** Comparison of the relative fitness of a Δ*mrrA* and a Δ*sigT* mutant under different conditions, each represented by an individual dot (30) (http://fit.genomics.lbl.gov/cgi-bin/compareGenes.cgi?orgId=Caulo&locus1=CCNA_03110&locus2=CCNA_03589). **(B)** Numbers and size of holdfast-mediated rosette structures are increased in a Δ*mrrA* mutant. Rosettes were analyzed in randomly chosen samples of exponentially growing cultures of wild type and the Δ*mrrA* mutant. **(C)** Comparison of c-di-GMP concentrations in *C. crescentus* wild type and a Δ*mrrA* mutant strain as determined by mass spectrometry analysis (see materials and methods).

**Figure S2:**
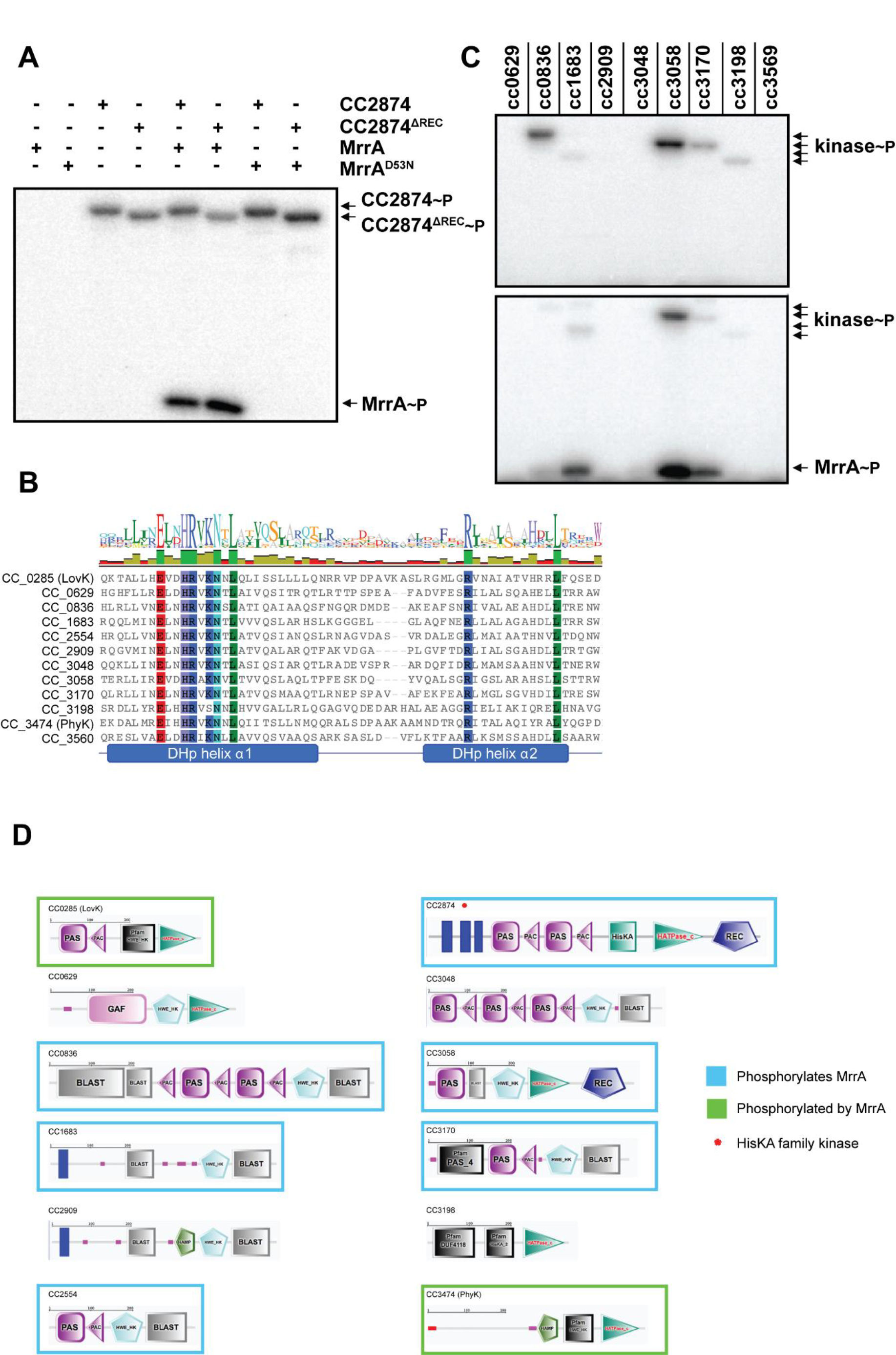
**(A)** Asp53 is the phospho-acceptor residue of MrrA. Autophosphorylation of histidine kinases CC2874 and CC2874 lacking its C-terminal receiver domain (ΔREC) and phospho-transfer to MrrA was analyzed in the presence of 500 µM ATP and 2.5 µCi [γ^32^P]ATP (3,000 Ci mmol^-1^). MrrA wild type or D53N mutant were used as indicated. Reactions were carried out for 15 minutes at room temperature and analyzed by SDS-PAGE and autoradiography. The position of phosphorylated proteins on the gel is indicated on the right. **(B)** Alignment of all *C. crescentus* HWE and HisKA2 histidine kinases based on the presence of the highly conserved HRXXN motif in their DHp domains (34). MUSCLE alignments were performed in Geneious v7.1.7 (www.geneious.com). **(C)** Autophosphorylation of nine *C. crescentus* HWE histidine kinases (upper panel) and phosphotransfer to MrrA (lower panel). Phosphorylation reactions were carried out as indicated in (A). **(D)** Schematic representation of the domain organization of the *C. crescentus* HWE/HisKA2 subfamily of histidine kinases. Highlighted with a red asterisk is the only HisKA kinase that phosphorylates MrrA. Graphics were generated by SMART (http://smart.embl-heidelberg.de).

**Figure S3:**
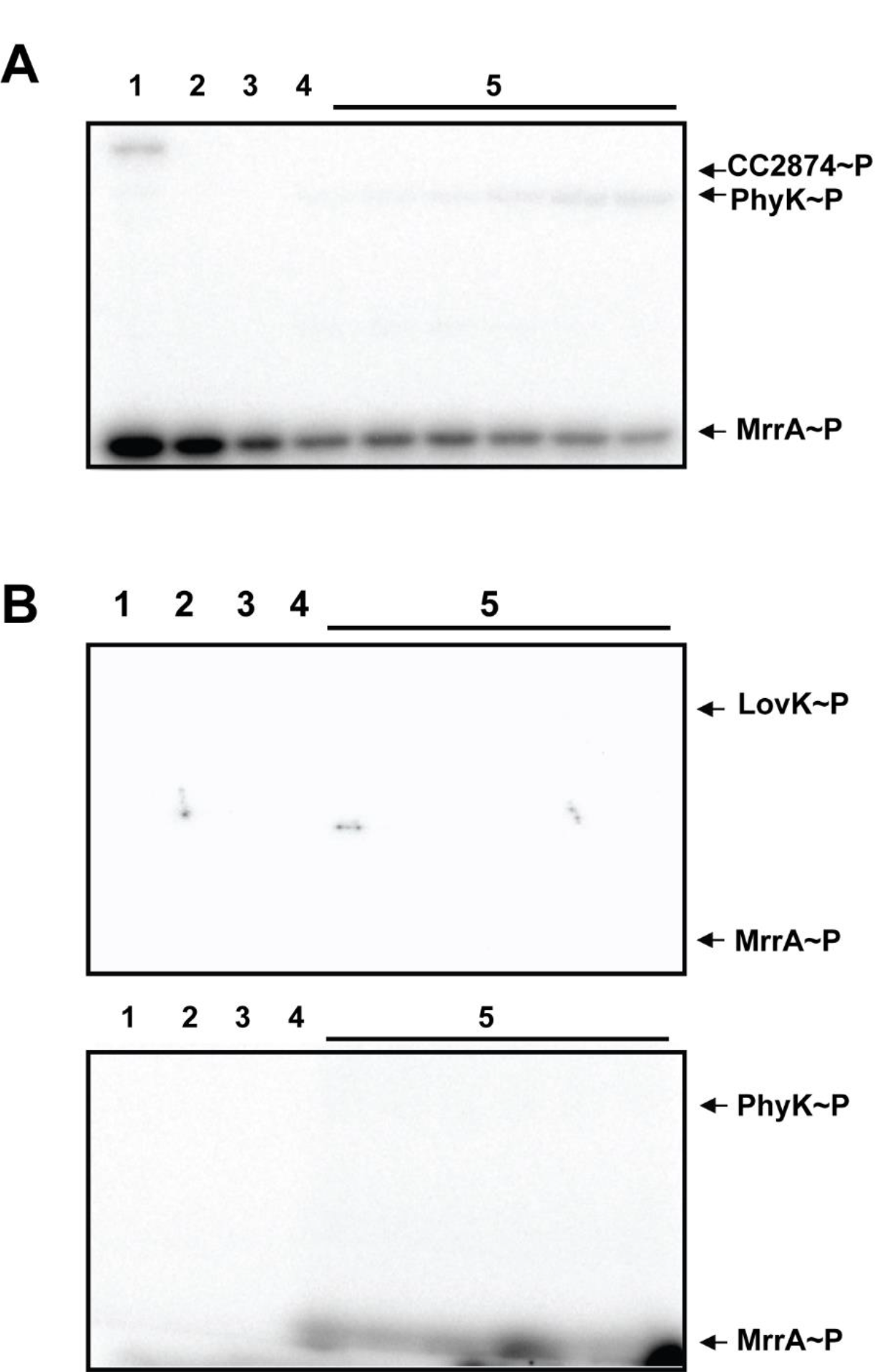
MrrA~P efficiently transfers phosphate to LovK and PhyK. **(A)**. Phosphorylation reactions with purified CC2874 and MrrA were carried out as in Figure 4F. The CC2874 kinase was then purified away using anti-MBP magnetic beads (lane 2). Remaining ATP was degraded using glucose and hexokinase (lane 3). Purified LovK was added with [γ^32^P]ATP and accumulation of LovK~P was monitored after 10 sec, 20 sec, 1 min, 2 min, 10 min and 20 min (lanes 5). Reactions were analyzed by SDS-PAGE and autoradiography. The position of phosphorylated proteins on the gel is indicated on the right. **(B)** Phosphorylation reactions as in (A) but MrrA was initially phosphorylated by CC2874 in the presence of unlabeled ATP (lane 1). Samples were then treated as in (A) (lanes 2-4) and purified LovK (upper panel) or PhyK (lower panel) was added together with [γ^32^P]ATP. Phosphorylation of LovK and PhyK was monitored over time as in (A).

